# Human Cytomegalovirus in breast milk is associated with milk composition, the infant gut microbiome, and infant growth

**DOI:** 10.1101/2023.07.19.549370

**Authors:** Kelsey E. Johnson, Timothy Heisel, David A. Fields, Elvira Isganaitis, Katherine M. Jacobs, Dan Knights, Eric F. Lock, Michael C. Rudolph, Cheryl A. Gale, Mark R. Schleiss, Frank W. Albert, Ellen W. Demerath, Ran Blekhman

**Affiliations:** Department of Genetics, Cell Biology, and Development, University of Minnesota, Minneapolis, USA; Division of Neonatology, Department of Pediatrics, University of Minnesota Medical School, Minneapolis, MN, USA; Department of Pediatrics, Diabetes-Endocrinology, University of Oklahoma Health Sciences Center, Oklahoma City, OK, USA; Pediatric, Adolescent and Young Adult Unit, Joslin Diabetes Center, Harvard Medical School, Boston, MA, USA; Department of Obstetrics, Gynecology and Women’s Health, Division of Maternal-Fetal Medicine, University of Minnesota Medical School, Minneapolis, MN, USA; BioTechnology Institute, College of Biological Sciences, University of Minnesota, Minneapolis, MN, USA; Department of Computer Science and Engineering, University of Minnesota, Minneapolis, MN, USA; Division of Biostatistics, University of Minnesota School of Public Health, Minneapolis, MN, USA; Harold Hamm Diabetes Center, Department of Physiology, Oklahoma University Health Sciences Center, Oklahoma City, OK, USA; Division of Pediatric Infectious Diseases and Immunology, University of Minnesota Medical School, Minneapolis, MN, USA; Division of Epidemiology and Community Health, University of Minnesota School of Public Health, Minneapolis, MN, USA; Section of Genetic Medicine, Division of Biological Sciences, University of Chicago, Chicago, IL, USA

**Author notes:** These authors jointly supervised the work (Frank W. Albert, Ellen W. Demerath, Ran Blekhman).

## Abstract

Human cytomegalovirus (CMV) is a highly prevalent herpesvirus that is often transmitted to the neonate via breast milk. Postnatal CMV transmission can have negative health consequences for preterm and immunocompromised infants, but any effects on healthy term infants are thought to be benign. Furthermore, the impact of CMV on the composition of the hundreds of bioactive factors in human milk has not been tested. Here, we utilize a cohort of exclusively breastfeeding full term mother-infant pairs to test for differences in the milk transcriptome and metabolome associated with CMV, and the impact of CMV in breast milk on the infant gut microbiome and infant growth. We find upregulation of the indoleamine 2,3-dioxygenase (IDO) tryptophan-to-kynurenine metabolic pathway in CMV+ milk samples, and that CMV+ milk is associated with decreased *Bifidobacterium* in the infant gut. Our data indicate a complex relationship between milk CMV, milk kynurenine, and infant growth; with kynurenine positively correlated, and CMV viral load negatively correlated, with infant weight-for-length at 1 month of age. These results suggest CMV transmission, CMV-related changes in milk composition, or both may be modulators of full term infant development.

## Introduction

Human Cytomegalovirus (CMV) is a member of the herpesvirus family with a global seroprevalence of ∼85% in women of childbearing age^1^. CMV is a double-stranded DNA virus that can infect multiple cell types including epithelial, endothelial, and immune cells^2^. Initial infection in healthy individuals is often asymptomatic, followed by lifelong viral latency. The most common mode of CMV transmission in infants is through breast milk, as during lactation CMV locally reactivates in the mammary gland in virtually all seropositive women^3–6^. Following mammary CMV reactivation, the presence of viral DNA in milk can be detected in both milk cells and whey^7–9^.

Postnatal CMV transmission via breast milk is thought to be benign in full-term, non-immunocompromised infants^10^. However, for preterm infants, postnatal CMV can have serious clinical consequences including sepsis, thrombocytopenia, and long-term neurodevelopmental impairment^10^. Among preterm and very low birthweight infants fed CMV+ breast milk, about 20% are estimated to acquire CMV^10,11^. The rate of transmission in full-term infants breastfed by seropositive mothers is estimated at up to 70%^12–14^.

Despite the prevalence and clinical importance of mammary CMV reactivation, little is known about its relationship to human milk composition. CMV reactivation could lead to a local immune response and viral regulation of host metabolism that could impact milk composition. Conversely, differences in milk composition could modify the risk of CMV reactivation, also leading to associations between CMV reactivation and milk composition. Associations between mammary CMV reactivation and the hundreds of nutritive and bioactive components of human milk have mostly not been assessed, but one study found an increase in pro-inflammatory cytokines in the setting of maternal CMV reactivation during lactation^5^. If CMV reactivation does alter human milk composition, it would be important to understand the impact of these changes on the infant. Variation in milk composition is associated with infant development, including the gut microbiome and immune system^15–17^. For preterm infants, who strongly benefit from human milk feeding^18^, an understanding of CMV-related changes in milk composition and their impact on infant health outcomes is critical.

One approach to understanding the mechanism by which CMV affects host physiology is to quantify the host transcriptional response and the metabolome in the context of CMV infection. The impact of CMV on host gene expression has been examined in cultured cells^19–24^ and in the blood of kidney transplant recipients^21^, but not in the context of mammary reactivation. Similarly, the metabolome during CMV infection has been described in cultured cells^25,26^ and infant urine^27^, but not in milk. Milk transcriptome and metabolome provide complementary profiles of the physiology of the lactating mammary gland and milk composition^15,28–30^. Although the clinical impact of postnatal CMV transmission is far greater for preterm than for term infants, the mechanisms by which CMV alters or is altered by human milk composition can be studied using milk from term mother-infant dyads.

In this study, we aimed to identify differences in human milk composition and infant outcomes associated with CMV reactivation in a deeply phenotyped cohort of lactating mothers and their full term infants.

Leveraging multi-omics data from mother-infant dyads, we tested for differences in the milk transcriptome, milk metabolome, and infant fecal metagenome associated with milk CMV reactivation (**Figure 1**). Further, we utilized anthropometric data to characterize differences in infant growth associated with milk CMV reactivation. Our results indicate that there are previously unappreciated differences in milk composition, infant gut microbiome composition, and growth in healthy full-term infants exposed to CMV through breast milk.

**Figure 1.**
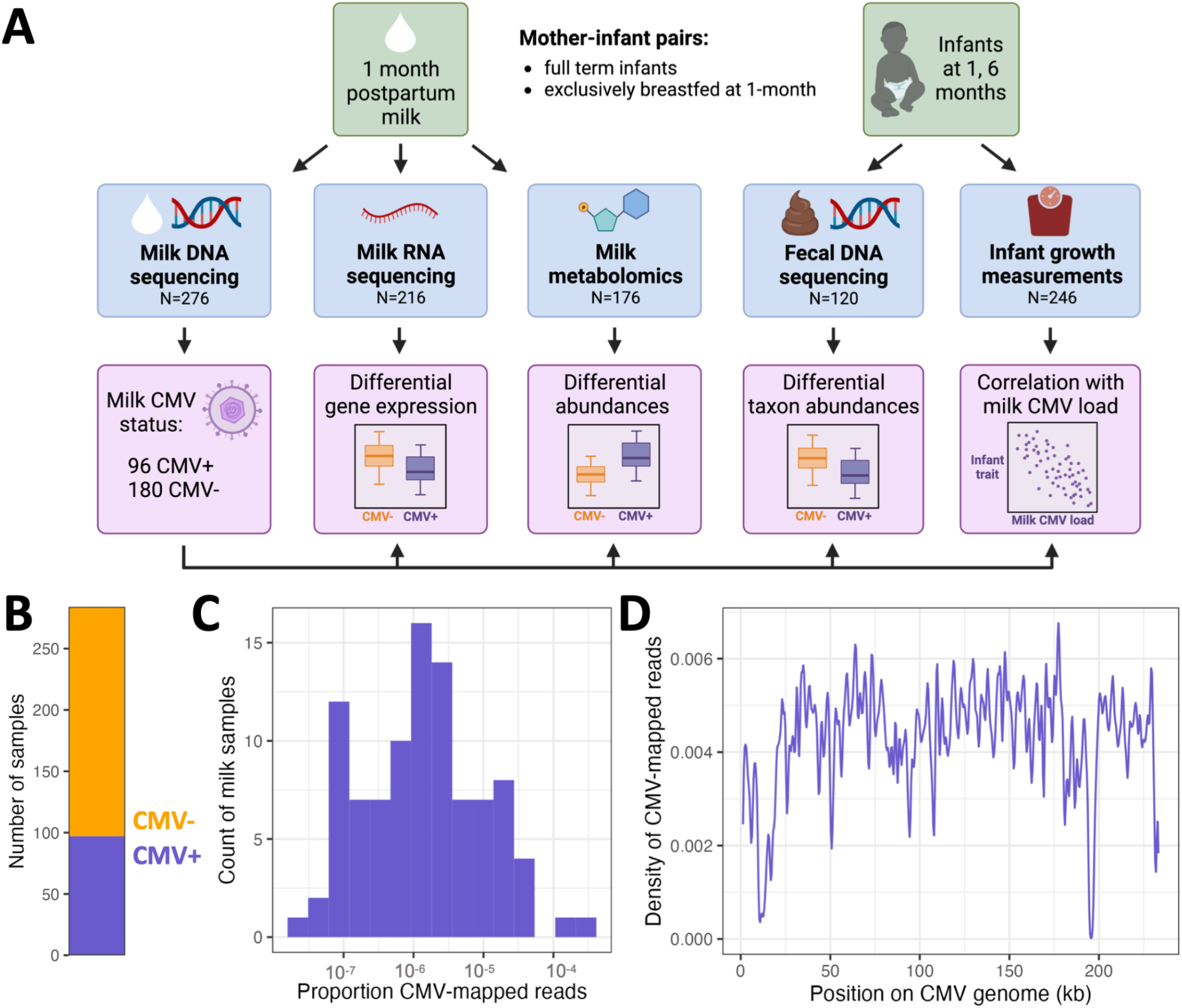
(A) Study overview. **(B)** Count of milk samples identified as CMV+ (N=97, purple) or CMV-(N=187, orange). **(C)** The distribution of CMV-mapped DNA reads, as a proportion of all DNA reads, across milk samples that had at least one read mapped to the CMV genome. **(D)** Density of CMV-aligned reads across the CMV genome from all CMV+ milk samples. The density refers to the fraction of all CMV-mapped reads aligned to a particular region of the CMV genome. The density dips close to zero at repetitive regions in the CMV genome^31^.

## Results

### Identifying CMV-positive samples from shotgun DNA sequencing of human milk

As CMV is a DNA virus, its presence can be detected in the lactating mammary gland by measuring CMV DNA in milk^8^. Viral shedding into breast milk typically begins within one week postpartum, and peaks 1-2 months postpartum^5^. We leveraged existing shotgun DNA sequencing data from 1-month postpartum milk samples^15^ (N=276) to identify milk samples with CMV viral shedding (**Figure 1A**). We mapped milk-derived DNA sequencing reads to the CMV genome and designated any sample with at least one read mapped to the CMV genome as CMV+ (97/284, 34% CMV+; **Figure 1B, Table S1**). Hereafter, samples with no CMV-mapped reads were designated as CMV-. To ensure our results were not dependent on this choice of threshold, we repeated the main analyses in this manuscript using a series of higher thresholds for the required proportion of CMV-mapped reads to designate a sample as CMV+. We saw no qualitative difference in our results across the range of tested thresholds (**Table S2**; but see infant growth section below). The mean proportion of CMV-mapped reads in samples designated as CMV+ was 1.0×10^−5^, or about 1 per 100,000 sequenced reads (**Figure 1C**), reflecting the fact that the vast majority of DNA in these milk samples comes from human cells.

Milk DNA was extracted and sequenced using two approaches for two distinct original goals: low-pass human whole genome sequencing (WGS) or shotgun metagenomic sequencing (SMS). The main difference between these approaches was the extraction protocol (see details in Methods). Within samples that had CMV-mapped reads from both datasets (N=24), there was a positive correlation in the proportion of CMV-mapped reads (Spearman’s rho=0.81, P=3.47×10^−5^; **Figure S1**). Mapped reads were widely distributed across the CMV genome (**Figure 1D**). There was no significant difference in the mean total read count for CMV+ vs. CMV-samples (two-sided t-test, P=0.26; **Figure S2**), suggesting that read depth did not bias our approach to detect CMV+ samples. Within CMV+ samples, there was no significant difference in the mean proportion of reads that mapped to the CMV genome between the two sources of DNA sequencing data (two-sided t-test, P=0.23; **Figure S3**). Taken together, these results suggest that our detection of CMV+ samples is not biased by technical factors or sequencing approach.

Comparing the maternal characteristics of CMV+ vs. CMV-milk samples, we observed that CMV+ milk samples were less likely to come from mothers who self-identified as White/European-American (74% in CMV+ vs. 91% in CMV-, P=3.1×10^−4^, q-value=3.7×10^−3^, Fisher’s exact test; **Table S3**). This is consistent with previous epidemiological estimates that CMV seropositivity is higher in non-white than white populations worldwide^32–34^. All other tested maternal traits were not significantly different between CMV+ and CMV-groups (q-value>0.25 for all other tests; **Table S3**).

### Immune response genes are upregulated in CMV+ milk samples

Human milk contains RNA from the milk-producing mammary epithelial cells and immune cells^35–38^. Thus, gene expression analyses of human milk provide a profile of the lactating mammary gland^28,29^. Using RNA-sequencing data we previously generated from the same milk samples studied here (N=221)^15^, we tested for differential expression of 17,675 genes in CMV+ vs. CMV-milk samples (**Figure 2A**). 36 genes were significantly differentially expressed (q-value<0.05), 34 of which were upregulated in CMV+ milk (**Figure 2B, Table S4**). These 34 upregulated genes were enriched for pathways related to the immune response to viral infections (**Table S5**), with “cellular response to interferon-gamma” as the most significant pathway (GO:0071346, odds ratio = 74.5, P = 5.22×10^−15^, q-value = 2.70×10^−12^). Upregulation of interferon-stimulated genes is a typical feature of the immune response to CMV infection^22,39,40^ (**Figure S4**). Within CMV+ milk samples, the proportion of CMV-mapped DNA reads and expression of the differentially expressed genes was significantly positively correlated for two genes: *BATF2* and *IDO1* (q-value<0.05, **Table S6**).

**Figure 2.**
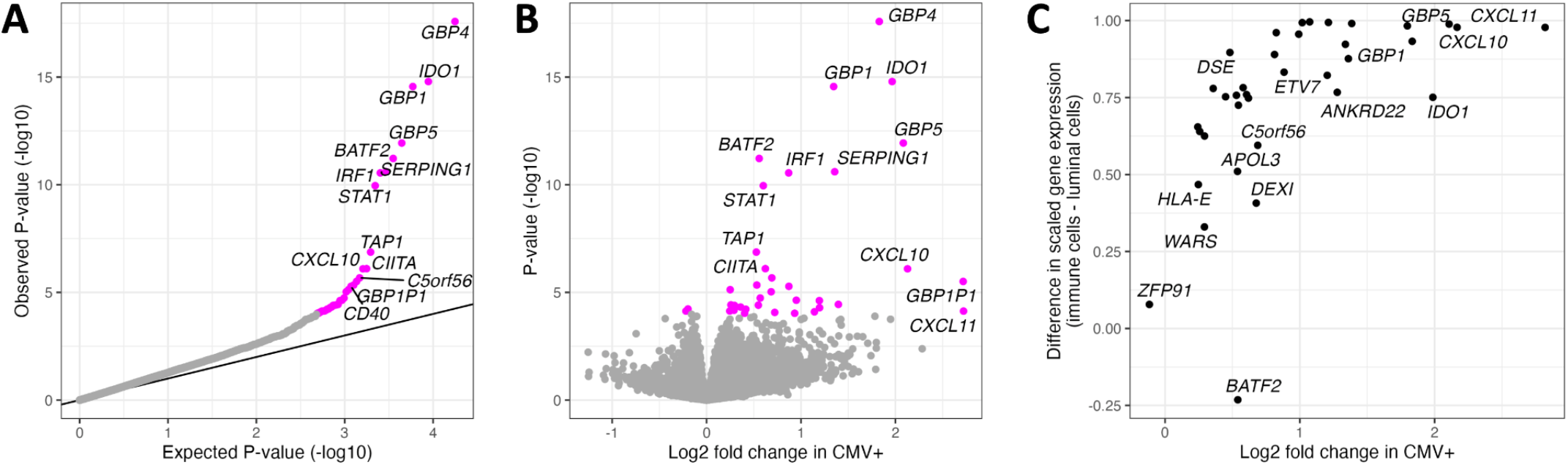
Differential gene expression analysis comparing CMV-to CMV+ milk samples. **(A)** QQ-plot from the results of differential gene expression analysis. The x-axis plots the expected P-value for the number of genes tested following a uniform distribution of P-values from 0 to 1, and the y-axis plots the observed P-values. Genes whose P-value was below the false discovery rate threshold of 5% are colored in magenta. **(B)** Volcano plot comparing estimated effect sizes of CMV+ on milk gene expression (x-axis) with each gene’s P-value (y-axis). Genes whose P-value was below the false discovery rate threshold of 5% are colored in magenta. **(C)** Comparison of log fold change in CMV+ samples from our bulk RNA-seq data (x-axis) vs. gene expression in a publicly available human milk single cell RNA-seq dataset^36^ (y-axis). Gene expression from milk single cells is plotted as the difference between scaled gene expression in immune cells and mammary luminal cells, to display that genes more highly expressed in our CMV+ milk samples tended to be more highly expressed in the immune cells in milk.

As our bulk milk RNA sequencing data derives from all the cells in our milk samples, we leveraged publicly available single cell RNA-sequencing data from human milk^36^ to explore the expression patterns of the 36 differentially expressed genes across milk cell types. We observed that genes more highly expressed in our CMV+ milk samples tended to also be more highly expressed in immune cells in milk in the single cell data (Spearman’s rho= 0.72, P= 1.7×10^−6^; **Figure 2C**). CMV+ milk samples also had a higher estimated proportion of immune cells (mean 16.5% in CMV+ vs. 12.6% in CMV-, P=0.041, Wilcoxon rank sum test; **Figure S5**; see Methods). We note that the CMV status of the milk samples in the reference single cell dataset (N=15) is unknown, but given the high prevalence of CMV it likely includes both CMV+ and CMV-samples. These results suggest that the elevated expression of these genes in CMV+ milk samples stems from an increased proportion of immune cells in CMV+ milk. This is potentially consistent with previous studies showing an increase in T cells in CMV+ human milk^40,41^, though we only tested the estimated proportion of all immune cells here due to the imprecision of cell-type deconvolution of bulk RNA-seq data.

### Differentially abundant metabolites in CMV+ samples indicate higher activity of the IDO tryptophan-to-kynurenine metabolic pathway

The human milk metabolome reflects cellular processes in the mammary gland and the composition of nutritive and bioactive components delivered to the infant^42^. We tested for differential abundance of 458 metabolites between 58 CMV+ and 84 CMV-milk samples in a regression model including study site, parity, maternal age, maternal pre-pregnancy BMI, maternal self-identified race, maternal gestational diabetes status, and maternal Healthy Eating Index score as covariates (**Figure S6**, see Methods). Two metabolites were significantly differentially abundant after correcting for multiple tests (q-value<0.05, **Table S7**): kynurenine (CMV+ estimated effect = 0.81, P= 1.3×10^−6^, q-value= 6.1×10^−4^; **Figure 3A**) and its metabolite kynurenic acid (CMV+ estimated effect = 0.75, P= 1.6×10^−5^, q-value= 6.6×10^−3^; **Figure S7A**).

**Figure 3.**
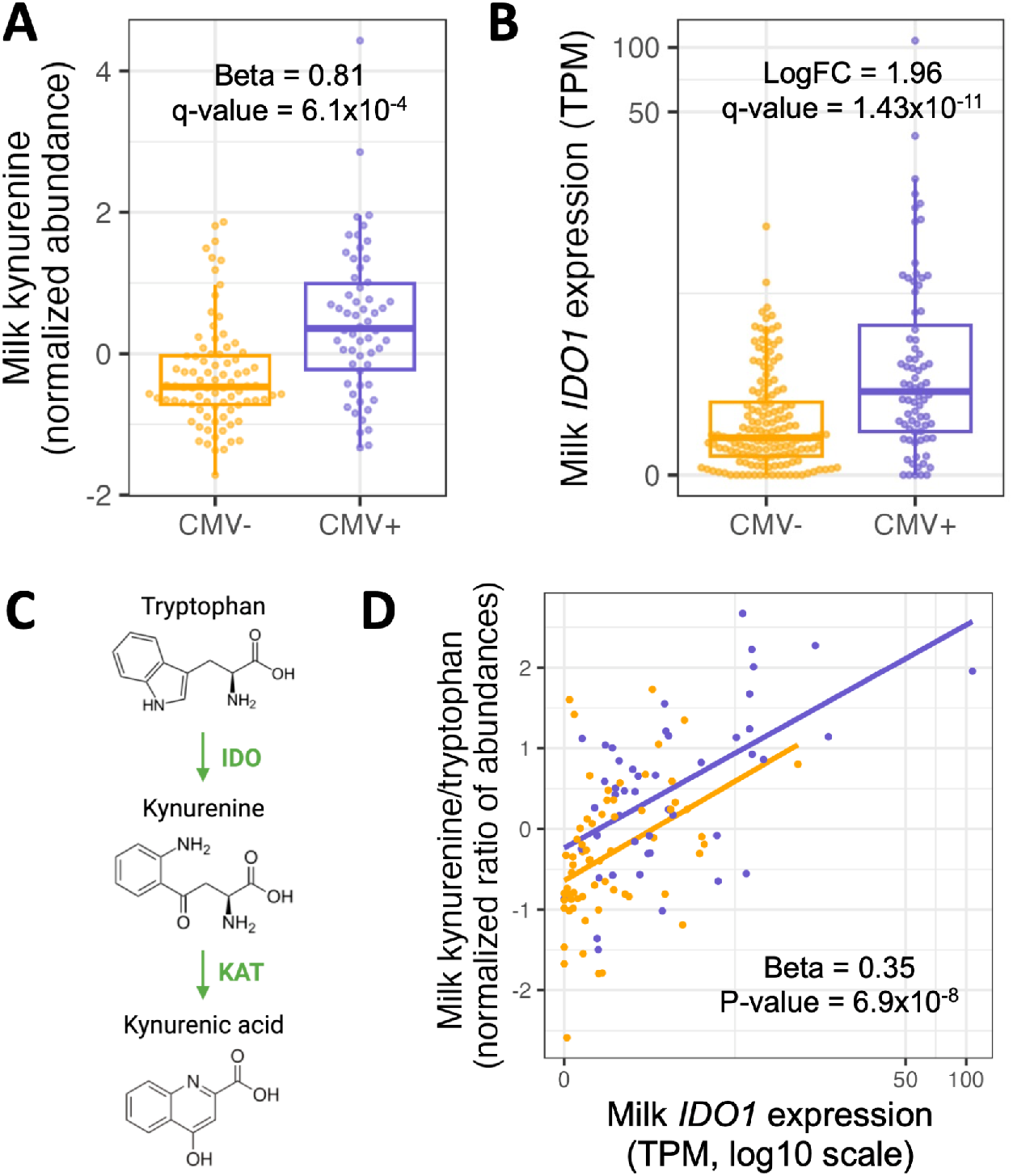
(A) Kynurenine abundances in CMV- (orange) vs. CMV+ (purple) milk samples. Each dot represents a milk sample. Plotted kynurenine levels (y-axis) are residuals after correcting for covariates included in the differential abundance analysis (see Methods). **(B)** *IDO1* expression in CMV-(orange) vs. CMV+ (purple) milk samples. Each dot represents a milk sample. LogFC: log fold-change between CMV+ and CMV-samples. **(C)** *IDO1* encodes the enzyme indoleamine 2,3-dioxygenase (IDO), which performs the rate-limiting step converting tryptophan to kynurenine. Kynurenic acid is metabolized from kynurenine by the KAT enzyme. **(D)** Correlation between *IDO1* expression (x-axis) and the ratio of kynurenine and tryptophan abundances (y-axis) in milk samples, stratified by CMV status. Each dot represents a milk sample.

The increased abundance of kynurenine and kynurenic acid in CMV+ samples is concordant with the upregulation of the *IDO1* gene we observed in our gene expression data (**Figure 3B**). *IDO1* encodes indoleamine 2,3-dioxygenase (IDO), the rate-limiting enzyme in the tryptophan-to-kynurenine metabolic pathway (**Figure 3C**). The kynurenine/tryptophan ratio was more significantly associated with CMV status than kynurenine alone (CMV+ estimated effect = 0.82, P= 9.4×10^−7^; **Figure S7B**). Within CMV+ milk samples, the kynurenine/tryptophan ratio was positively correlated with the proportion of CMV-mapped reads (Beta = 0.19, P = 6.3×10^−3^; **Figure S7C**). We did not observe a difference in the abundance of tryptophan by CMV status (CMV+ estimated effect = -0.22, P= 0.20, q-value= 0.85). Milk *IDO1* expression was also positively correlated with the kynurenine/tryptophan ratio of abundances in milk, independent of milk CMV status (Beta= 0.35, P=6.9×10^−8^; **Figure 3D**), illustrating the strong link between expression of *IDO1* and the abundance of these metabolites.

### Milk CMV status is correlated with composition of the infant gut microbiome

Variation in human milk composition has been previously associated with the infant gut microbiome^15,43,44^. Motivated by the differences in milk composition we observed between CMV+ and CMV-milk samples, we next tested for associations between milk CMV status and composition of the infant gut microbiome. We previously generated shotgun metagenomic data from infant feces collected at 1 and 6 months postpartum (N=127 mother/infant pairs at 1 month, N=120 at 6 months)^15,45^. To explore a potential relationship between milk CMV status and the overall structure of the infant fecal microbiome, we first reduced the dimensionality of the microbial taxon abundance table using principal component analysis (each time point analyzed separately, see Methods). We then tested for associations between milk CMV status and the microbial principal components (PCs). Milk CMV status was significantly correlated with PC3 of the 1-month infant fecal metagenomes (Beta=1.79, P=1.1×10^−3^, q-value = 5.6×10^−3^; **Figure 4A, Table S8**). The top-loading taxa in 1-month PC3 were species of *Bifidobacterium* (negatively correlated with PC3; **Figure S8**). PC3 was not correlated with milk kynurenine abundance (P=0.12); and within infants fed CMV+ milk, PC3 was not correlated with the proportion of CMV-mapped reads in milk (P=0.79). Milk CMV status was not associated with the 6-month taxon abundance PCs (**Table S8**). Separately, we performed principal component analysis on the microbial genetic pathway abundances estimated from shotgun metagenomic data, and milk CMV status was not associated with any of the pathway PCs (**Table S9**). Milk CMV status was not associated with infant fecal alpha diversity at 1 month (Beta=0.29, P=0.15) or 6 months (Beta=0.06, P=0.70).

**Figure 4.**
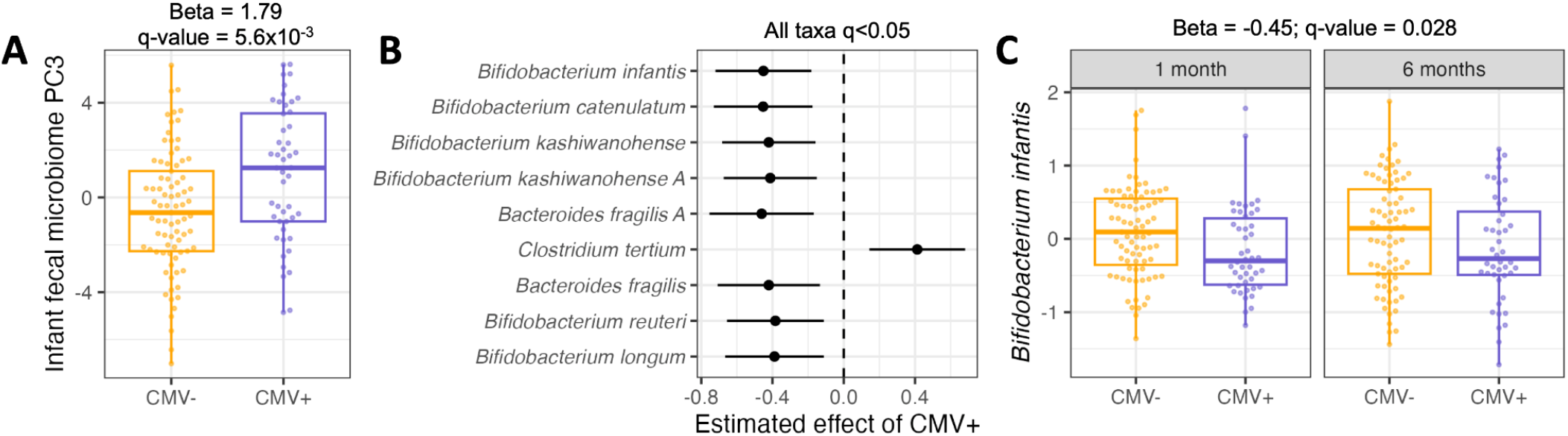
(A) Comparison of PC3 values for infant fecal samples fed CMV- (orange) vs. CMV+ (purple) breastmilk. Principal component analysis was performed on the taxon abundance table for infant fecal samples at 1 month of age. Each dot represents an infant fecal sample. Plotted PC3 levels are residuals after correcting for covariates included in the association analysis with milk CMV status (see Methods). **(B)** Estimated effect of CMV+ milk on normalized microbial taxa abundances in the infant gut, including samples from 1 and 6 months of age. All taxa listed had a P-value below a false discovery rate of 5%. Taxa are arranged from smallest (top) to largest (bottom) P-value. **(C)** The distribution of *Bifidobacterium infantis* abundances in the infant fecal microbiome, for infants fed CMV- (orange) or CMV+ (purple) milk, at 1 and 6 months of age. Plotted *B. infantis* levels are residuals after correcting for covariates included in the association analysis with milk CMV status (see Methods). In **(B)** and **(C)**, taxon relative abundances were centered log ratio transformed and scaled to mean 0, standard deviation 1 before association analysis.

We next tested for associations between milk CMV status and abundances of individual microbial taxa. We modeled 56 microbial species’ abundances in both 1 and 6 month old infants in a linear mixed effects model (see Methods). Abundances of nine taxa were significantly correlated with milk CMV status (q-value<0.05), including six species of *Bifidobacterium* that were less abundant in the gut metagenomes of infants fed CMV+ milk; *Clostridium tertium*, which was more abundant in infants fed CMV+ milk; and *Bacteroides fragilis*, which was less abundant in infants fed CMV+ milk (**Figure 4B, Table S10**). The taxon with the strongest association with milk CMV status was *Bifidobacterium infantis* (Beta = -0.45, P = 1.4×10^−3^, q-value = 0.028; **Figure 4C**).

### Milk CMV status is correlated with infant growth

Finally, we tested if exposure to CMV+ milk was associated with infant growth, measured as weight-for-length Z-score (WLZ), a commonly used nutritional status metric to assess adequacy of weight relative to length and age in infants^46^. Infants fed CMV+ milk had on average approximately one-third of a Z-score greater weight-for-length at 1 month of age compared to infants fed CMV-milk (Beta = 0.38, P=0.011, N=246; linear regression including WLZ at birth and additional covariates, see Methods; **Figure 5A, Table S11**). This relationship between WLZ and 1 month milk CMV status was not present at birth or at 6 months of age (**Figure 5A, Figure S9A**). Infants fed CMV+ milk had somewhat lower mean length-for-age Z-score at 1 month (Beta = -0.27, P=0.025, **Figure 5A, Figure S9B**), and no difference in weight-for-age Z-score at 1 month (Beta = -0.012, P=0.89, **Figure 5A, Figure S9C**). These results indicate that infants fed CMV+ milk in the first month of life tended to have weight growth that exceeded their length growth in the first month. However, this difference did not persist to 6 months of age.

**Figure 5.**
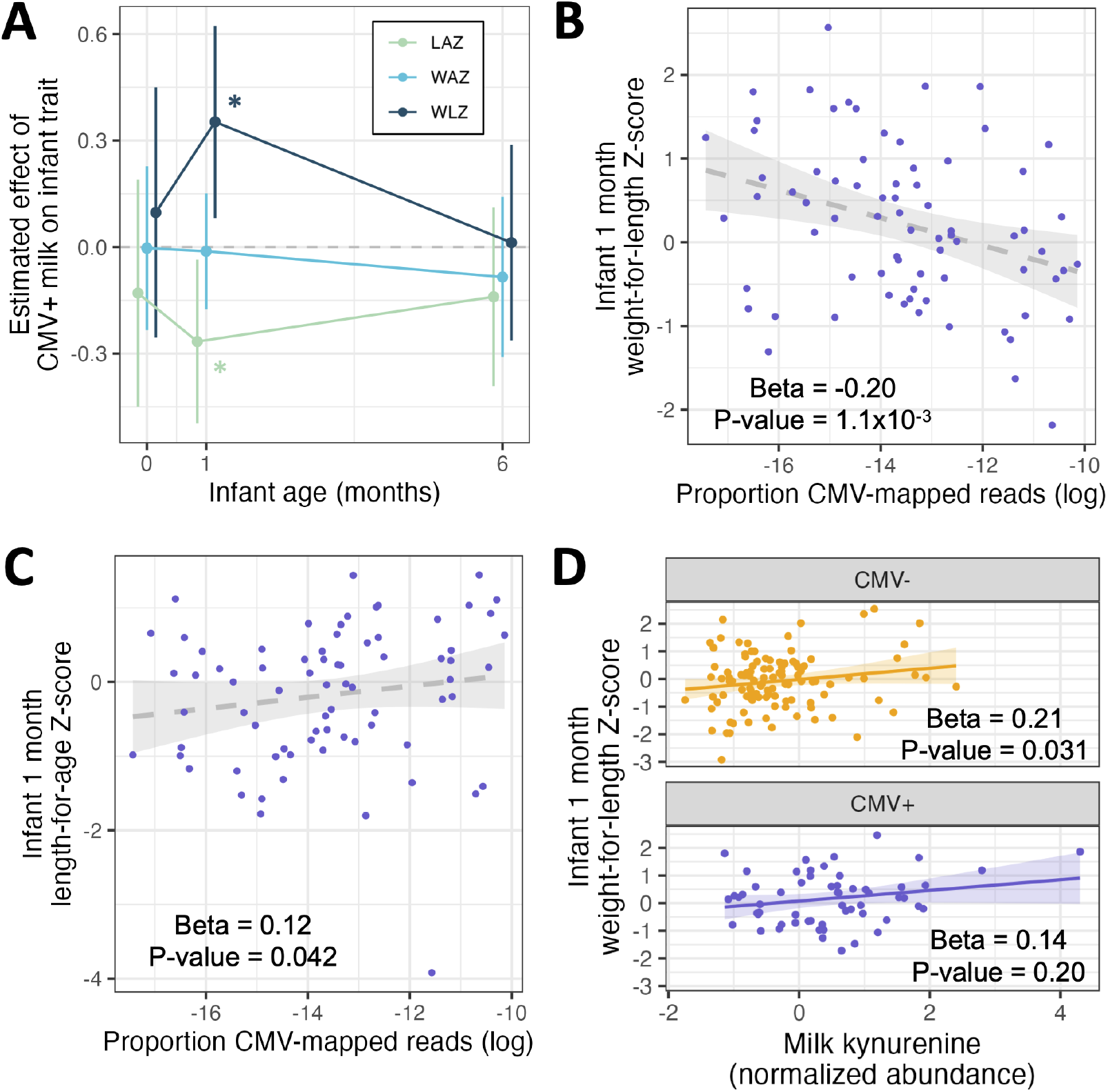
Results of multivariate regressions of infant anthropometric measurements vs. milk CMV status, proportion CMV-mapped reads in milk, or milk kynurenine. All regression models included the equivalent Z-score at birth as a covariate (except when the Z-score at birth was the response variable). **(A)** Estimated effect of CMV+ milk on infant growth metrics at birth, 1 month, and 6 months of age. Error bars represent 95% confidence intervals. *P<0.05; LAZ: length-for-age Z-score; WAZ: weight-for-age Z-score; WLZ: weight-for-length Z-score. **(B)** Within infants fed CMV+ milk, there was a negative correlation between the proportion of CMV-mapped reads and infant WLZ at 1 month of age. **(C)** Within infants fed CMV+ milk, there was a positive correlation between the proportion of CMV-mapped reads and infant WLZ at 1 month of age. **(D)** There was a positive correlation between the abundance of kynurenine in milk and infant WLZ at 1 month, when tested for infants fed CMV+ (orange, top) or CMV-(purple, bottom) milk separately. All plotted infant growth metrics in panels B-D are residuals after correcting for covariates included in the association analyses with milk CMV status (see Methods).

Within infants fed CMV+ milk, we observed a negative correlation between the proportion of CMV-mapped reads in milk and infant WLZ at 1 month (Beta = -0.20, P = 1.1×10^−3^, N=74; **Figure 5B**), the opposite direction of the relationship when comparing CMV- and CMV+ groups. We also observed a positive correlation between the CMV-mapped read proportion in milk and infant length-for-age at one month (Beta = 0.12, P = 0.042; **Figure 5C**), and no correlation with infant weight-for-age at one month (Beta = -0.035, P = 0.46; **Figure S10**). The relationship between milk CMV load and infant growth can be seen in our sensitivity analysis of escalating thresholds to designate milk samples as CMV+: as the threshold increases, only the milk samples with the highest proportion of CMV-mapped reads are designated CMV+, and the effect estimate of CMV+ milk on infant WLZ and milk CMV status reverses direction from positive to negative (**Table S2**). These results suggest that a factor other than CMV viral load itself is driving the CMV group differences in WLZ at 1 month.

Hypothesizing that the relationship between CMV status and infant growth could be due to CMV-related differences in milk composition, we tested for a relationship between milk kynurenine abundance and infant 1-month WLZ. Kynurenine was positively correlated with WLZ (Beta = 0.21, P = 1.9×10^−3^; **Figure S11**), a relationship that persisted when milk CMV status was added as a covariate (Beta = 0.20, P = 0.011). Further, when testing the relationship between milk kynurenine and infant WLZ in CMV+ and CMV-groups separately, there was a positive correlation for both groups; though, it was only significant in the CMV-group (CMV+: Beta = 0.14, P = 0.20; CMV-: Beta = 0.24, P = 0.031; **Figure 5D**). Within infants fed CMV+ milk, when including both milk kynurenine and the proportion of CMV-mapped reads in milk, both terms were correlated with infant WLZ in opposing directions (kynurenine: beta = 0.22, P = 0.047; CMV read proportion: beta = -0.15, P = 0.011). Given that (1) accounting for milk kynurenine levels removes the association between milk CMV status and infant WLZ at one month; and (2) CMV viral load is correlated with WLZ in the opposite direction as milk CMV status, even when including milk kynurenine levels; we conclude that increased kynurenine in CMV+ milk samples, or a correlated factor, is responsible for the positive association between milk CMV status and infant WLZ at 1 month.

## Discussion

In this study, we found that the presence of CMV DNA in human milk is associated with milk gene expression and metabolite abundances, altered composition of the infant gut microbiome, and potential disruptions to infant growth in the first month of life. Notably, our study utilized a cohort of healthy, full-term infants in the U.S.; a population where the impact of CMV presence in breast milk or postnatal CMV transmission was largely thought to be negligible.

We utilized shotgun DNA sequencing from the cell pellet of human milk to identify samples with the presence of CMV at 1 month postpartum. Our study demonstrates that non targeted DNA sequencing of human milk can be used to identify CMV+ samples. We identified CMV DNA in 32% of 1 month milk samples, which is lower than the estimated prevalence for US adults of childbearing age (∼50%)^32,47^. Given that virtually all seropositive women will have CMV reactivation in the mammary gland during lactation^48^, and CMV viral loads are estimated to peak around 4-6 weeks postpartum^5,8^, we likely were unable to detect CMV in some samples with a low viral load. We also acknowledge that while viral reactivation during lactation is likely the primary cause of CMV DNA in breast milk, CMV could also be shed through breast milk in the context of primary infections or re-infections occurring late in gestation.

Using complementary milk RNA sequencing and metabolomics approaches, we identified an upregulation of the *IDO1* tryptophan-to-kynurenine metabolic pathway in CMV+ milk samples. This pathway has previously been implicated in the immune response to CMV in studies of human cells and primary tissues^49,50^, suggesting this association may be a response to mammary CMV reactivation. Additionally, one study found that providing kynurenine to human fibroblasts promoted CMV replication, and blocking *IDO1* decreased CMV replication^51^. Given our observational study design, we cannot determine if the association with increased *IDO1*/kynurenine is a cause or consequence of mammary CMV reactivation. Overall, the impact of CMV on milk composition was notably narrow, with a handful of genes and two metabolites differentially abundant between CMV+ and CMV-milk samples.

Under conditions of chronic viral infection, activation of the IDO pathway can lead to a more tolerogenic immune state^49^, but the impact of elevated milk kynurenine and its metabolites on the infant is unknown. Kynurenine induction of the aryl hydrocarbon receptor (AHR) can cause immunosuppression via generation of regulatory T-cells^52^, and AHR activation may protect against necrotizing enterocolitis and inflammation in the infant gut^53,54^. Whether kynurenine metabolites in milk are at high enough concentrations to have physiological effects in the infant, and the potential impacts of CMV on this pathway, are possible areas of future investigation. We observed a positive association between milk kynurenine and infant growth at 1 month, with higher milk kynurenine correlated with lower length-for-age and greater weight-for-length Z-scores, suggesting milk kynurenine levels could impact growth in early life independent of CMV status. It is important to note that while the impact of kynurenine on weight-for-length was of moderate effect statistically, differences of this magnitude are not generally of clinical significance for healthy term infants.

We also observed that within infants fed CMV+ milk, higher CMV-mapped read proportion (as a proxy for viral load) was negatively correlated with infant weight-for-length and positively correlated with length-for-age at 1 month of age. Previous research on the impact of postnatal CMV transmission on infant growth has primarily focused on two contexts: (1) very low birth weight infants in the NICU setting, and (2) in perinatally HIV-exposed but uninfected infants. Studies focused on very low birth weight infants have found mixed evidence for impacts of postnatal CMV on anthropometric measures^55^. The largest study to date in very low birth weight infants found that postnatal CMV acquisition was associated with lower weight-for-age Z-score at discharge, but no difference in length-for-age in a U.S. population^56^. In HIV-exposed but uninfected Malawian infants, breast milk CMV DNA load was negatively correlated with infant weight-for-length, length-for-age, and weight-for-age at 6 months (infant CMV status was unknown in this study)^57^. In addition, a study of Zambian infants found that postnatal CMV acquisition was associated with lower length-for-age Z-score at 18 months in both HIV-exposed and HIV-unexposed infants, but no difference in weight-for-age by CMV status^58^. The context of our cohort is quite different from these previous analyses, yet cumulatively, these studies suggest that postnatal exposure and/or acquisition of CMV can impact infant growth.

We observed that exposure to CMV+ milk was associated with the composition of the infant gut microbiome in our cohort of breastfed babies. Specifically, CMV+ milk-exposed infants had lower abundances of *Bifidobacterium* species and higher abundances of *Clostridium tertium*. Lower *Bifidobacterium*, particularly *B. infantis*, in the infant gut microbiome is associated with adverse health outcomes^59–61^. *C. tertium* has been reported as potentially pathogenic in the infant gut^62,63^. Notably, milk kynurenine was not associated with the infant gut microbiome in our study, indicating that the potential effects of CMV viral load on infant growth and the infant gut microbiome may act through distinct pathways. A previous study by Sbihi et al. examined the impact of CMV acquisition on the infant gut microbiome. In a population-based birth cohort, early CMV acquisition (in the first 3 months of life) but not later CMV acquisition (between 3-12 months) was associated with lower alpha diversity (i.e. within-sample diversity)^64^. While our study is not directly comparable as we do not know infant CMV status, we did not observe a significant difference in alpha diversity in CMV-exposed vs. unexposed infants. Sbihi et al. also observed increased incidence of childhood allergy with early CMV acquisition^64^, a phenotype not currently assessed in our cohort.

A limitation of our study is the unknown serostatus of the infants at birth and subsequent timepoints, as infant blood samples were not available. As previous studies estimate up to 70% of breastfed term babies of seropositive mothers acquire CMV postnatally^12–14^, it is possible that a substantial fraction of the babies fed CMV+ milk in our study had postnatally acquired CMV by 1 month of age. The infants in our study were also not tested for congenital CMV, which has a prevalence of about 4.5 per 1000 births in the US^65,66^ and is often asymptomatic and undetected^67^. Further studies are required to characterize the impacts of CMV+ milk on growth and the gut microbiome in infants with and without CMV transmission.

While there is growing awareness and understanding of the negative impacts of breastmilk-acquired CMV in preterm infants^10^, it is generally thought to be benign in healthy term infants. Some have even speculated that there may be an evolutionary advantage to postnatal CMV acquisition, in the form of a ‘natural immunization’ or other immune-boosting effect for the infant^12^. We find that exposure to CMV+ milk is associated with reduction in beneficial microbes in the infant gut. Given the high prevalence of CMV globally, impacts on infant microbiome development could have a substantial impact at the population level. This study highlights not only these CMV-related changes but also more generally, how ‘normal’ variation in human milk impacts healthy infant development.

## Acknowledgements

We would like to thank Katy Duncan, Laurie Foster, Tipper Gallagher, and all MILk study staff and participants for their contributions, and members of the Albert and Blekhman labs for helpful discussions related to this project. This work was supported by the resources and staff at the University of Minnesota Genomics Center (https://genomics.umn.edu). This work was carried out in part by resources provided by the Minnesota Supercomputing Institute (https://www.msi.umn.edu/).

## Author contributions

Conceptualization: KEJ, CAG, MRS, FWA, EWD, RB

Formal analysis: KEJ, TH

Funding acquisition: KEJ, DK, KMJ, EFL, DAF, CAG, FWA, EWD, RB

Investigation: KEJ, TH

Supervision: MCR, MRS, CAG, FWA, EWD, RB

Writing - original draft: KEJ

Writing - review and editing: All authors

## Data availability

Milk metabolite abundances, gene expression matrices, and microbial abundance tables are available as extended data tables (see descriptions in supplementary material). Raw sequencing data will be available at dbGaP prior to publication.

## Materials and Methods

### Description of study population

This study made use of existing data from the Mothers and Infants LinKed for Healthy Growth (MILK) study. Recruitment protocols and study characteristics have previously been extensively described^15,45,68–71^. This study recruited mothers intending to exclusively breastfeed their infants prenatally. Study visits occurred at two sites: the University of Minnesota (MN) or the University of Oklahoma Health Sciences Center (OK). All included infants were born at full term. The milk samples utilized in this manuscript were collected during a study visit about 1 month postpartum via a full breast milk expression two hours after a complete infant feed. Expressed milk volume and weight was recorded, milk was gently mixed, aliquots were made, and then stored at -80°C within 20 minutes of collection and kept at -80°C until thawed for RNA/DNA extraction or metabolomics analysis.

### Human milk RNA extraction, sequencing, and gene expression quantification

The human milk RNA extraction protocol, sequencing, and gene expression quantifications used in this study have been previously described^15^. RNA extraction, library preparation, and sequencing was performed at the University of Minnesota Genomics Center (UMGC). Briefly, bulk RNA was extracted from the whole milk cell pellet to profile gene expression of all cell types present in the milk sample. RNA was extracted from the cell pellet using the RNeasy Plus Universal HTP following the manufacturer’s instructions. RNA libraries were prepared with the TakaraBio Stranded Total RNA Pico Mammalian kit and sequenced on an Illumina NovaSeq 6000 S2 flow cell with 2×150 paired-end reads in two pools. Gene-level quantifications were generated using RNA-SeQC v2.3.4^72^.

### Analyses with publicly available single cell RNA-seq data from human milk

Raw gene counts (MIT_Milk_Study_Raw_counts.txt.gz) and metadata (MIT_milk_study_metadata.csv.gz) were downloaded for the Nyquist et al. study^73^ from the Broad Insitute Single Cell Portal (https://singlecell.broadinstitute.org/single_cell/study/SCP1671/cellular-and-transcriptional-diversity-over-the-course-of-human-lactation) on 6/3/2022. Gene counts for each cell were scaled to log(*x*/*s* + 1), where *x* was the gene count in a cell and *s* was a scaling factor. *s* was calculated as the total counts per cell divided by the mean of total counts across all cells. For each of the 36 differentially expressed genes in our CMV+ milk samples, the scaled expression for each cell type was calculated as the mean scaled expression across all cells of that cell type, divided by the gene’s mean scaled expression in the cell type with the highest mean expression. In Figure 2C, immune cell expression included six cell types (T cells, eosinophils, dendritic cells, B cells, neutrophils, macrophages) and mammary luminal cell expression included two cell types (luminal cell 1 and luminal cell 2).

Cell type proportions were estimated for each milk sample with bulk RNA-sequencing data as previously described^15^, using a publicly available single cell RNA sequencing dataset from human milk^36^ cells and Bisque^74^. Proportions of 8 cell types were estimated: two types of mammary epithelial cells (luminal cell 1, luminal cell 2) and six immune cell types (T cells, eosinophils, dendritic cells, B cells, neutrophils, macrophages). The estimated immune cell proportion was calculated as the sum of the six immune cell types.

### Human milk DNA extraction and sequencing

DNA was extracted and sequenced from human milk using separate protocols for different initial applications

1. **Human low-pass whole genome sequencing (WGS)**: The DNA extraction protocol and sequencing for this application has been previously described^15^. In brief, DNA was extracted from the cell pellet at UMGC with the QIAamp 96 DNA Blood Kit, and sequenced by Gencove, Inc. for target sequencing depth of ∼1x for the human genome.
2. **Shotgun metagenomic sequencing (SMS)**: DNA extraction and sequencing from milk samples for this application has also been previously described^15,45^. DNA was extracted using the PowerSoil kit, libraries constructed for metagenomics sequencing using the Illumina Nextera XT kit, and sequenced on an Illumina NovaSeq system using the S4 flow cell with the 2×150 bp paired end V4 chemistry kit at UMGC.

### Identification of CMV-positive milk samples

We mapped DNA sequencing reads generated from human milk with the above two approaches to the human cytomegalovirus genome to identify milk samples with CMV DNA. Starting with the WGS DNA reads, we mapped the reads from each milk sample to seven CMV genome isolates from human milk ^75^ accessed from NCBI Genbank (https://www.ncbi.nlm.nih.gov/genbank/,MW528458 – MW528464) using Bowtie2^76^. Finding that the number of aligned reads across reference CMV isolates was in strong agreement, we continued with the aligned read count for each sample from isolate BM1 (accession MW528458) for reads from both WGS and SMS. We called milk samples as CMV+ if they had at least one concordantly mapped read pair with MAPQ>5 from either WGS or SMS. Of the 276 milk samples utilized in this study, 86 had both WGS and SMS (n=34 CMV+), 132 only had WGS (n=40 CMV+), and 58 had only SMS (n=22 CMV+). The proportion of CMV-mapped reads was calculated for each CMV+ sample as the number of reads mapped to the CMV genome divided by the total number of sequencing reads, with counts from SMS and WGS data summed if both were available.

### Identification of differentially expressed genes by milk CMV status

Differential gene expression analysis between CMV- and CMV+ milk samples was performed in DESeq2^77^ using the gene-level read count matrix generated with RNA-SeQC^72^. 17,675 genes were included in differential gene expression analysis. Maternal age, maternal pre-pregnancy BMI, maternal self-reported race, maternal parity, infant age in days, sample RIN, RNA sequencing pool, and the mass RNA extracted from the sample were included as covariates. None of the individuals with transcriptomic data had gestational diabetes, so this was not included as a covariate. P-values were adjusted for multiple tests using the default Benjamini and Hochberg method in DESeq2^77,78^. Enrichment analysis of upregulated genes was performed with EnrichR^79^, using “GO_Biological_Process_2021” as the reference gene ontology. To test for a correlation between CMV-mapped read proportion and gene expression, the same DESeq2 model was used, replacing CMV status with the CMV-mapped read proportion (logged and scaled to mean 0, s.d. 1) and including only CMV+ samples.

### Human milk metabolomics and identification of differentially abundant metabolites

Samples for milk metabolomics were prepared and analyzed as previously described^80^ from frozen milk samples at BERG health (Framingham, MA). For each of 458 metabolites, the association between metabolite abundance and milk CMV status was estimated using a multivariate regression with ‘lm’ in R. Metabolite abundances were log(x+1) transformed and scaled to mean 0, standard deviation 1. Additional included covariates were the study center (MN vs. OK), parity, maternal age, maternal pre-pregnancy BMI, maternal gestational diabetes (yes/no), maternal self-reported race (white vs. non-white) and maternal Healthy Eating Index total score^81^ (averaged from three timepoints: prenatal, 1 month postpartum, and 3 months postpartum). P-values were corrected for multiple tests using the Benjamini-Hochberg false discovery rate^78^ with ‘p.adjust’ in R. To test for a correlation between CMV-mapped read proportion and metabolite abundance, the same multivariate model was used, replacing CMV status with the CMV-mapped read proportion (logged and scaled to mean 0, s.d. 1) and including only CMV+ samples.

### Infant fecal metagenomics and comparison with milk CMV status

Infant fecal sample collection, DNA extraction, metagenomic sequencing, and estimation of microbial taxon and pathway abundances from 1 and 6 month samples has been previously described^15,45^. Principal components analysis of 1 and 6 month infant metagenomes, summarized as taxon or pathway abundances, was performed separately. Data were filtered to include only taxa/pathways with relative abundance >0.001 in at least 10% of 1-month or 6-month samples. A centered log-ratio transformation was performed on the relative abundances of each sample, and principal components were calculated with the ‘prcomp’ command in R. Associations between the metagenomic PCs that explained at least 5% of the variance in the data (5 PCs each for 1 and 6 month taxa abundances, 3 PCs each for pathway abundances at 1 and 6 months) and milk CMV status were calculated using linear regression with the ‘glm’ command in R. Infant delivery mode (cesarean vs. vaginal), maternal parity, maternal age, maternal self-identified race, maternal pre-pregnancy BMI, maternal gestational diabetes (yes/no), maternal Group B streptococcus status, fecal sample collection site (home vs. study visit), and maternal Healthy Eating Index total score^81^ (averaged from three timepoints: prenatal, 1 month postpartum, and 3 months postpartum) were included as covariates. Two additional covariates were included in the regression models for 6 month infant fecal samples: exclusive breastfeeding status at 6 months (yes/no), and if complementary foods had been introduced at 6 months (yes/no). At 1 month, all infants were exclusively breastfed with no complementary foods. Additional variables about antibiotics use were not included (beyond Group B Streptococcus status, which is treated with antibiotics during labor) because there was too much missing data that would vastly reduce the sample size for these analyses.

Alpha diversity was calculated for each infant fecal sample from 1 or 6 months with the inverse Simpson index with the unfiltered taxon count matrix using the vegan^82^ package in R. Alpha diversity was scaled to mean 0, s.d. 1 and tested for association with milk CMV status in a multivariate regression model including the same covariates described above for the microbiome PCs.

Associations between individual taxon abundances and milk CMV status were estimated using a linear mixed effects model with the ‘lmerTest’ package^83^ in R. Using taxon abundances (centered log-transformed and scaled to mean 0, standard deviation 1 within each timepoint) from both 1 and 6 month timepoints as the response variable; fixed effects variables were milk CMV status, sample time point (1 or 6 months, coded as 0 or 1), infant delivery mode (cesarean or vaginal), maternal parity, maternal self-reported race, maternal pre-pregnancy BMI, maternal Group B streptococcus status, fecal sample collection site (home vs. study visit), maternal gestational diabetes (yes/no), and exclusive breastfeeding status at 6 months; and the mother-infant pair ID was included as a random variable. Only species-level taxa with relative abundance >0.001 in at least 10% of samples in both 1 and 6 month samples were included (56 species). P-values were corrected using the Benjamini-Hochberg false discovery rate with ‘p.adjust’ in R.

### Infant growth measurement and comparison with milk CMV status

Infant growth measurements and Z-score calculation from this cohort have been previously described^70,84^. Age and sex-specific length-for-age, weight-for-age, and weight-for-length Z-scores (WLZ) were calculated using the World Health Organization standards for term infants^46^. Association between infant 1-month WLZ and milk CMV status was calculated in a regression model including WLZ at birth, infant race (parental report), maternal pre-pregnancy BMI, maternal gestational diabetes (yes/no), household income, and delivery mode (cesarean vs. vaginal) as covariates with the ‘lm’ command in R. Associations between milk CMV status and 3- and 6-month WLZ were calculated in the same model, replacing the outcome (1-month WLZ) with the 3-or 6-month WLZ. Associations with length-for-age or weight-for age Z-scores used the same covariates, replacing WLZ at birth with the respective Z-score at birth. To test for a correlation between CMV-mapped read proportion and WLZ, CMV status was replaced in the model with the CMV-mapped read proportion (logged and scaled to mean 0, s.d. 1) and including only CMV+ samples.

