## Supplementary material for "Human Cytomegalovirus in breast milk is associated with milk composition, the infant gut microbiome, and infant growth"

K.E. Johnson, et al. 2023

**Figure S1.**

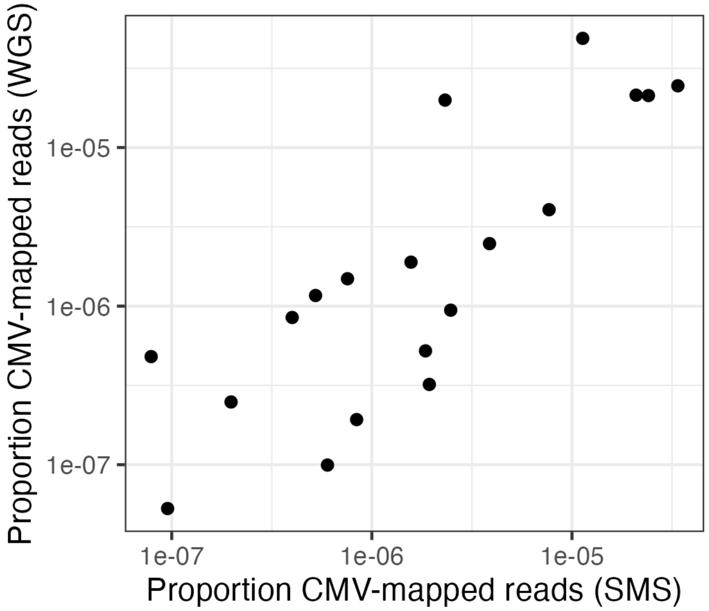

**Figure S1.** Correlation between the proportion of CMV-mapped reads for CMV+ milk samples with sequencing data from both sources used in this study (N=24, Spearman's rho=0.81, P=3.47x$10^{-5}$).

**Figure S2.**

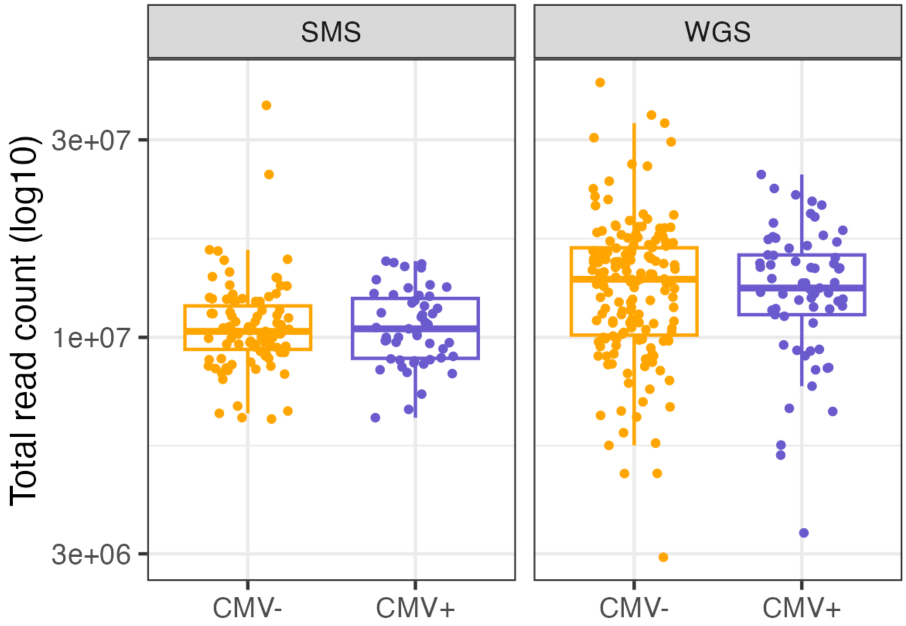

**Figure S2.** The distribution of total read counts for CMV+ vs. CMV- milk samples from either shotgun metagenomic sequencing (SMS) or whole genome sequencing (WGS). There was no significant difference in mean total read count between CMV- and CMV+ milk samples in either dataset.

**Figure S3.**

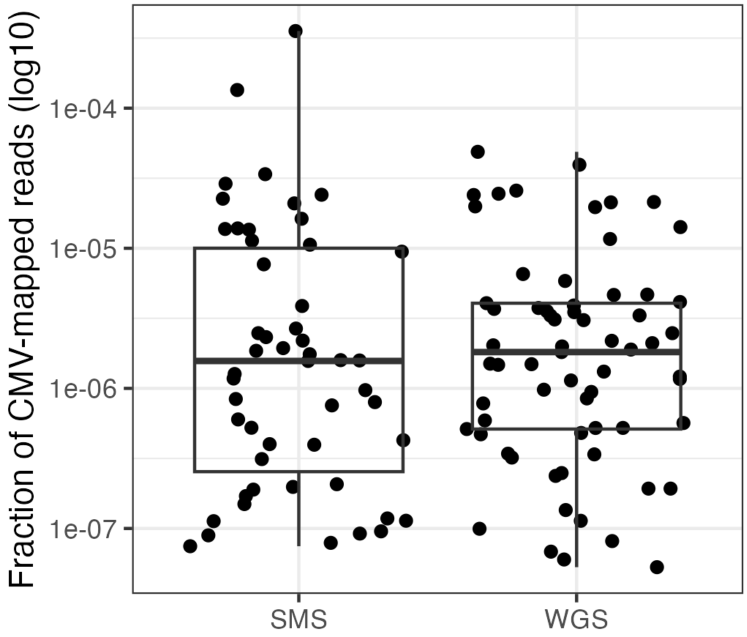

**Figure S3.** The distributions of the proportion of CMV-mapped reads for milk samples with either shotgun metagenomic sequencing (SMS) or whole genome sequencing (WGS). There was no significant difference in the mean proportion of CMV-mapped reads between the two data types.

**Figure S4.**

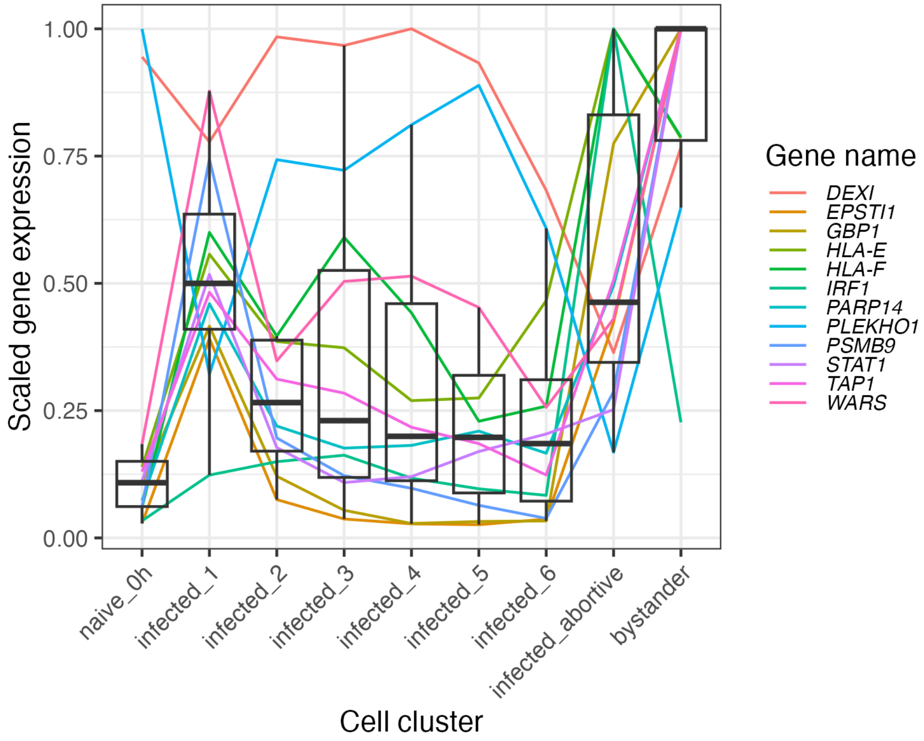

**Figure S4**. For 12 genes that were upregulated in our CMV+ milk samples, this plot shows expression patterns across cell clusters in a publicly available dataset of single cells (human fibroblasts) exposed to CMV (Hein & Weissman, 2022). The x-axis groups are the 9 cell type clusters identified in the single cell dataset; naive_0h: cells before CMV infection; infected_1, …, infected_6: cell clusters along the CMV infection trajectory; infected_abortive: cells who are initially infected by CMV but the infection does not proceed to viral replication; bystander: uninfected cells that have high expression of interferon response genes due to signaling from nearby infected cells. The y-axis is expression values, scaled relative to the expression level in the cell cluster where each gene was most highly expressed. 10 out of 12 genes were most highly expressed in the ‘bystander’ cluster, which were defined as cells that did not have viral gene expression but did express high levels of interferon response genes.

**Figure S5.**

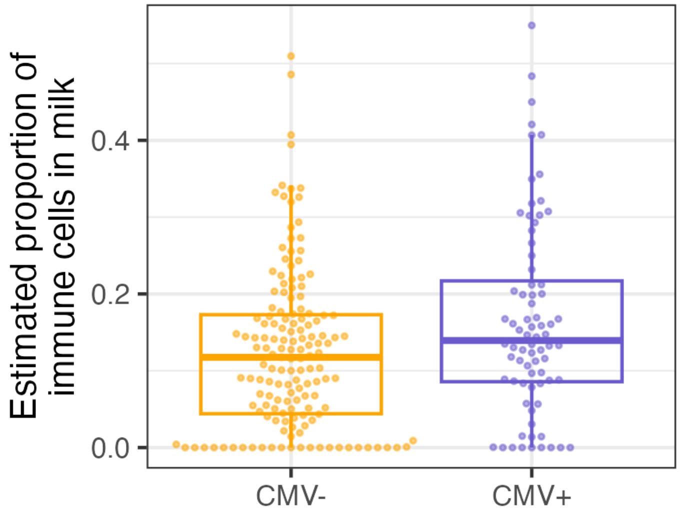

**Figure S5.** The estimated proportion of immune cells in milk for CMV+ (purple) vs. CMV- (orange) milk samples. Cell type proportions were estimated via deconvolution of bulk RNA-sequencing data using a publicly available reference human milk single cell RNA-sequencing dataset.

**Figure S6.**

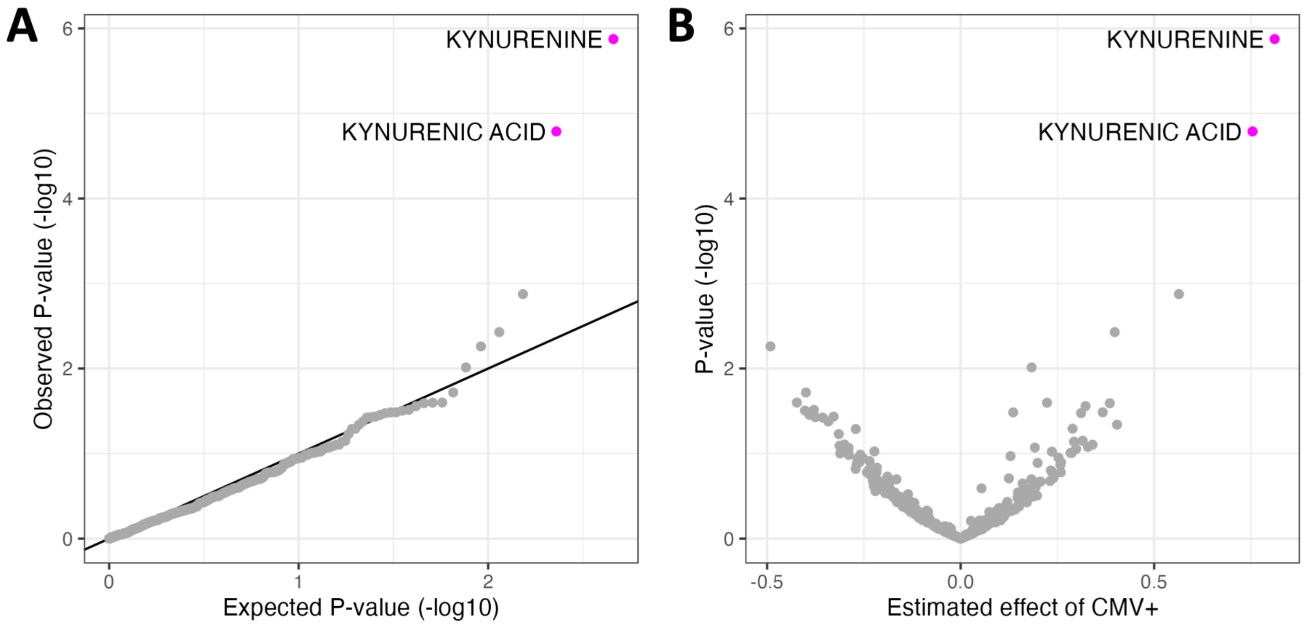

**Figure S6. (A)** QQ-plot from the results of differential abundance analysis comparing metabolites in CMV+ and CMV- milk samples. The x-axis plots the expected P-value for the number of metabolites tested following a uniform distribution of P-values from 0 to 1, and the y-axis plots the observed P-values. **(B)** A volcano plot showing estimated effect sizes of CMV+ on milk metabolite abundance (x-axis) with each metabolite's P-value (y-axis). Metabolites whose P-value was below the false discovery rate threshold of 5% are colored in magenta.

**Figure S7.**

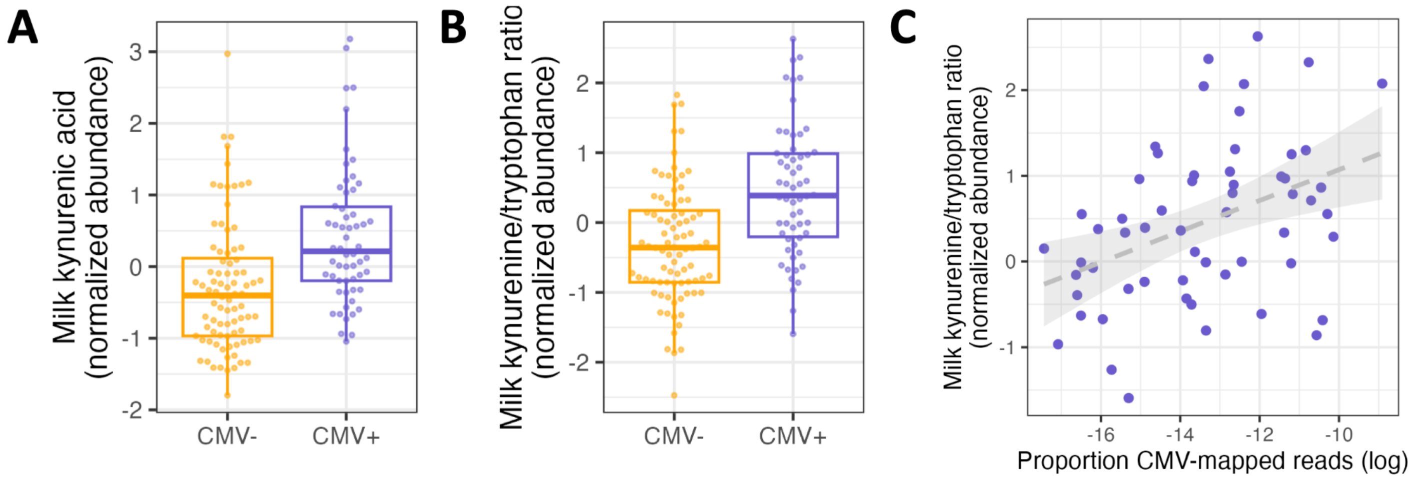

**Figure S7. (A)** Kynurenic acid abundances in CMV- (orange) vs. CMV+ (purple) milk samples (beta = 0.75, P= $1.6 \times 10^{-5}$, q-value= $6.6 \times 10^{-3}$). Each dot represents a milk sample. **(B)** CMV+ milk samples (purple) had a higher ratio of kynurenine/trytophan abundances compared to CMV- (orange) (beta = 0.83, P = $2.7 \times 10^{-6}$). Each dot represents a milk sample. **(C)** We observed a positive correlation between the proportion of CMV-mapped reads in each CMV+ milk sample (x-axis) and the ratio of kynurenine/tryptophan abundances (y-axis) (Beta = 0.19, P = $6.3 \times 10^{-3}$). Each dot represents a milk sample, and only CMV+ milk samples (purple) were included in this analysis. All plotted metabolite abundances are residuals after correcting for covariates included in the association analysis with milk CMV status (see Methods).

**Figure S8.**

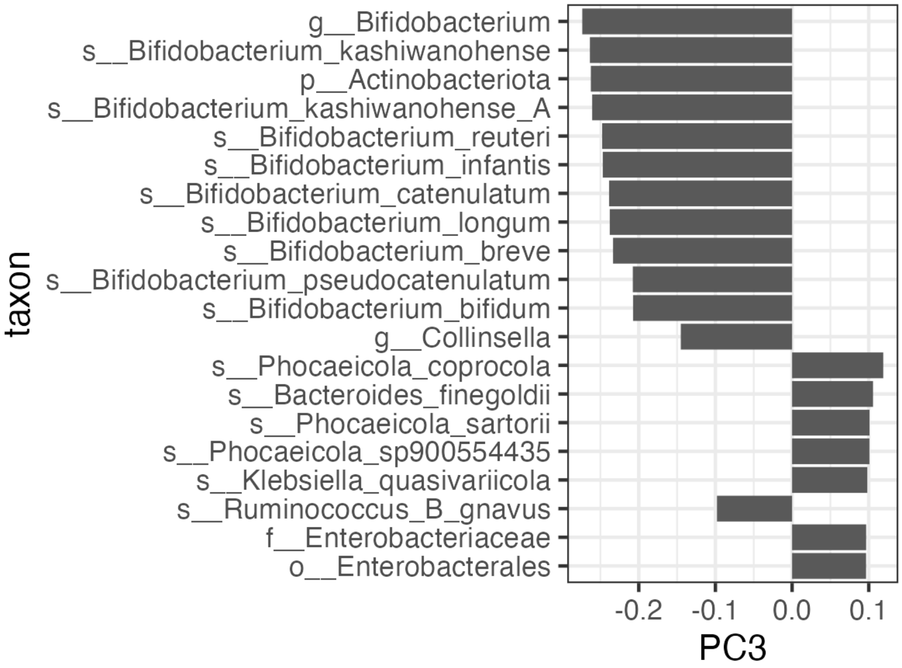

**Figure S8.** The top 20 loading taxa for PC3 of the 1 month infant fecal metagenomes. For each taxon, the bar magnitude and direction represents its loading on PC3. Taxa are sorted in order of greatest (top) to smallest (bottom) magnitude, with the top 20 taxa included in this plot.

**Figure S9.**

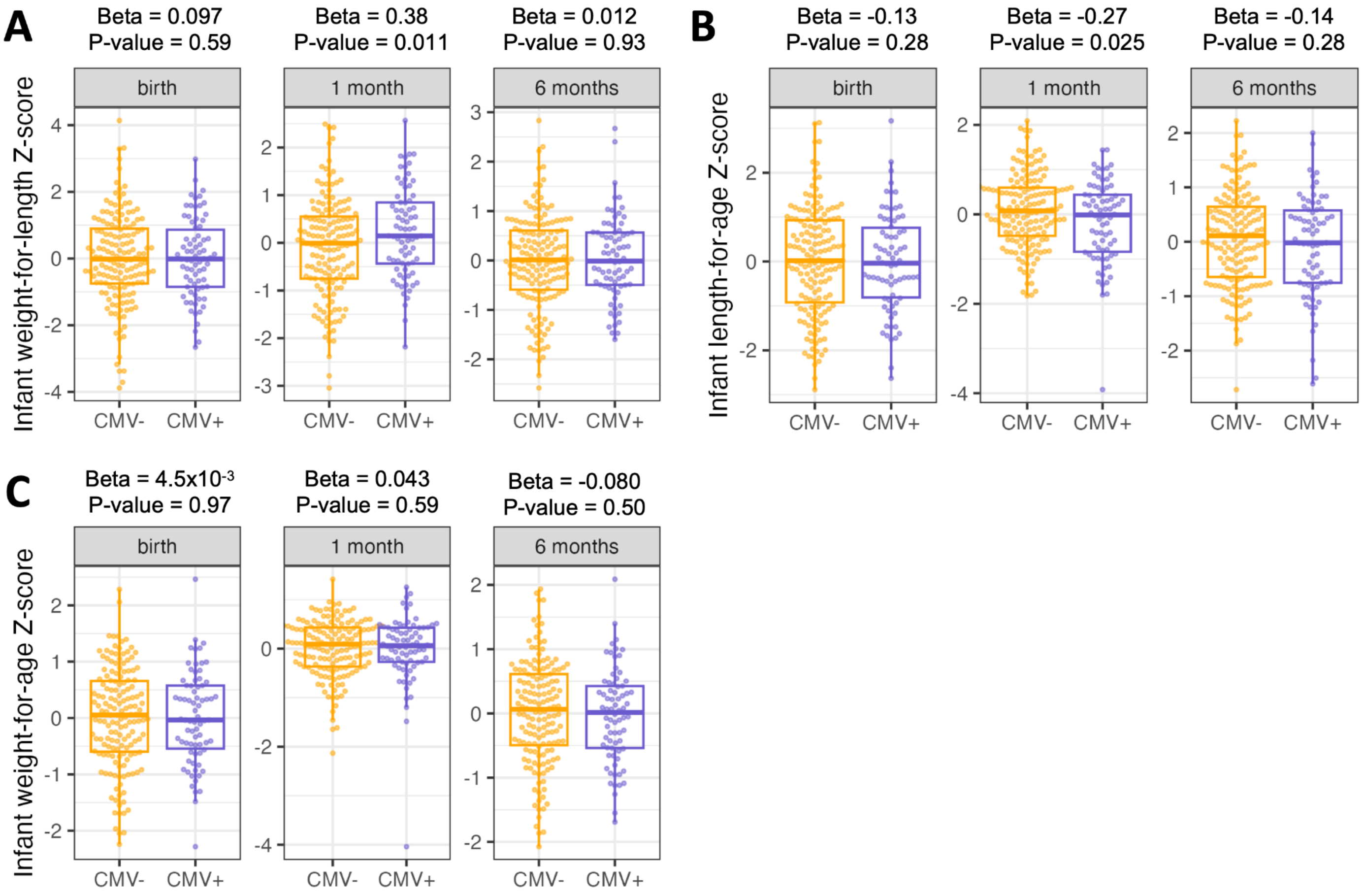

**Figure S9.** Results of multivariate regressions of infant weight-for-age Z-score at birth, 1 month of age, or 6 months of age vs. milk CMV status at 1 month postpartum. All regression models included the equivalent Z-score at birth as a covariate (except Z-score at birth). All plotted infant growth metrics are residuals after correcting for covariates included in the association analyses with milk CMV status (see Methods).

**Figure S10.**

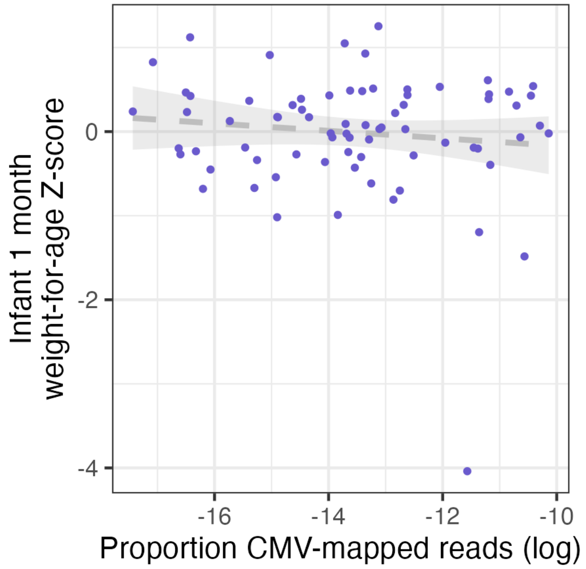

**Figure S10.** Within infants fed CMV+ milk, there was no correlation between the proportion of CMV-mapped reads and infant weight-for-age Z-score at 1 month of age (Beta = -0.035, P = 0.46). Plotted infant growth metrics are residuals after correcting for covariates included in the association analysis with milk CMV status (see Methods).

**Figure S11.**

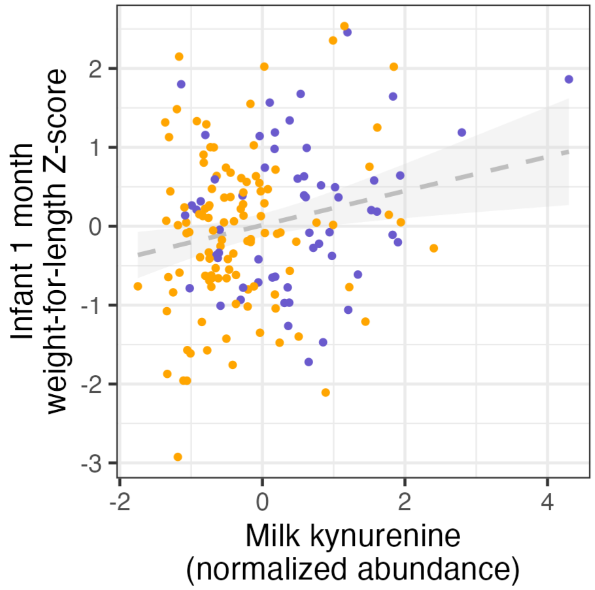

**Figure S11.** There was a positive correlation between milk kynurenine (x-axis) and infant 1-month WLZ (y-axis). Each point represents a mother/infant pair, CMV- milk in orange, CMV+ milk in purple. Plotted infant growth metrics are residuals after correcting for covariates included in the association analysis with milk CMV status (see Methods).

**Supplementary Table Descriptions**

**Table S1.** Milk sample CMV status data and other metadata. **indID**: mother/infant pair unique ID. **sms.cmv**: count of reads mapping to CMV genome from milk shotgun metagenomic sequencing data. **wgs.cmv**: count of reads mapping to CMV genome from milk whole genome sequencing data. **sms.tot**: total reads in milk shotgun metagenomic sequencing data. **wgs.tot**: total reads in milk whole genome sequencing data. **all.cmv**: total number of milk DNA sequencing reads mapping to CMV genome. **all.tot**: total number of milk DNA sequencing reads. **cmv.prop**: proportion of all milk DNA sequencing reads mapping to CMV genome. **milk.cmv.status**: milk sample designation as CMV+ or CMV-. **Center**: study site (0=MN, 1=OK). **Parity**: number of previous births. **matage**: maternal age. **mat_bmi**: maternal pre-pregnancy BMI. **delivery_cat**: delivery mode (0: vaginal, 1: cesarean). **mateduc_cat**: maternal education category (0=high school/GED/associate's, 1=bachelor's degree, 2=graduate degree). **income_cat**: household income category (0 = less than $30k, 1 = between $30k to $90k, 2 = greater than $90k). **wlz_0, wlz_1, wlz_6**: infant weight-for-length Z-score at birth, 1 month, or 6 months of age. **laz_0, laz_1, laz_6**: infant length-for-age Z-score at birth, 1 month, or 6 months of age. **waz_0, waz_1, waz_6**: infant weight-for-age Z-score at birth, 1 month, or 6 months of age. **gdm**: gestational diabetes status during the focal pregnancy. **mat.white**: maternal self-identify as white/European-American. **ebf6mo**: exclusive breastfeeding status at 6 months postpartum. **inf.white**: infant self-identify (by mother) as white/European-American. **ebf3mo**: exclusive breastfeeding status at 3 months postpartum. **avg_totalhei**: average total Healthy Eating Index score from surveys during pregnancy, 1 month postpartum, and 3 months postpartum. **compfoods6mo**: complementary solid foods introduced to infants diet at 6 months of age. **gbs**: maternal Group B streptococcus status at delivery. **fecal_1mo_site**: collection site of 1 month infant fecal sample (home vs. study visit). **fecal_6mo_site**: collection site of 6 month infant fecal sample (home vs. study visit).

**Table S2.** Sensitivity of main results to the proportion of CMV-mapped reads in milk used as a threshold to call a sample CMV+. Five thresholds were tested: the 1%, 10%, 25%, and 50% quantiles of the distribution of CMV-mapped read proportion within milk samples with at least 1 CMV-mapped read.

**Table S3**. Associations between milk CMV status and maternal traits. **CMV+, CMV-**: For binary traits, the percentage of participants in the CMV+/CMV- category with the trait is listed, with the number of participants with the trait listed in parentheses. For continuous traits, the mean trait value is given with the 2.5% and 97.5% percentiles in parentheses. **CMV+ N, CMV- N**: The number of participants in each CMV status group for each trait (some traits had missing data so the numbers were not the same for every trait). **P-value**: the P-value of the test for difference in proportions (binary traits, two-sided Fisher's exact test) or difference in mean trait value (continuous traits, two-sided t-test) between CMV+ and CMV- groups. **Q-value**: Benjamini-Hochberg corrected P-value. **Trait**: the tested trait.

**Table S4**. Output of DESeq2 testing for differential gene expression between CMV+ and CMV- milk samples. **baseMean**: the average of the normalized count values. **log2FoldChange**: the fold change (log2 scale) in CMV+ compared to CMV- milk samples. **lfcSE**: standard error of the log2 fold change. **stat**: value of the test statistic for significance testing. **pvalue**: P-value of significance testing for a differential gene expression between CMV+ and CMV- milk samples. **padj**: Adjusted P-value after Benjamini-Hochberg correction. **gene_id**: Ensembl gene ID. **gene_name**: Gene symbol.

**Table S5**. Output of pathway enrichment testing. **Term**: tested gene ontology. **Overlap**: number of overlapping genes, number of genes in pathway. **P.value**: Enrichment test P-value. **Adjusted.P.value**: Benjamini-Hochberg corrected P-value. **Odds.Ratio**: odds ratio of overlap. **Genes**: overlapping genes.

**Table S6**. Output of DESeq2 testing for correlation between gene expression and the proportion of CMV-mapped reads in milk samples. **baseMean**: the average of the normalized count values. **log2FoldChange**: the fold change (log2 scale) change in gene expression for each standard deviation change in CMV-mapped read proportion. **lfcSE**: standard error of the log2 fold change. **stat**: value of the test statistic for significance testing. **pvalue**: P-value of significance testing for a differential gene expression between CMV+ and CMV- milk samples. **padj**: Adjusted P-value after Benjamini-Hochberg correction. **gene_id**: Ensembl gene ID. **gene_name**: Gene symbol.

**Table S7**. Results of differential abundance testing of metabolites in CMV+ vs. CMV- milk samples. **beta**: estimated change in metabolite abundance in CMV+ compared to CMV- milk samples. **se**: standard error of effect estimate. **p**: P-value of effect estimate. **q**: Benjamini-Hochberg corrected P-value. **metabolite**: tested metabolite.

**Table S8**. Results of association testing between principal components of the 1 and 6 month infant fecal microbiome taxon abundances and milk CMV status. **beta**: estimated effect of CMV+ vs. CMV- milk on PC value. **se**: standard error of effect estimate. **p**: P-value of effect estimate. **q**: Benjamini-Hochberg corrected P-value. **PC**: principal component tested. **timepoint**: 1 or 6 month infant fecal samples.

**Table S9.** Results of association testing between principal components of the 1 and 6 month infant fecal microbial pathway abundances and milk CMV status. **beta**: estimated effect of CMV+ vs. CMV- milk on PC value. **se**: standard error of effect estimate. **p**: P-value of effect estimate. **q**: Benjamini-Hochberg corrected P-value. **PC**: principal component tested. **timepoint**: 1 or 6 month infant fecal samples.

**Table S10.** Results of association testing between milk CMV status and microbial species abundances in infant fecal samples at 1 and 6 months of age. **est**: estimated effect of CMV+ vs. CMV- milk on PC value. **se**: standard error of effect estimate. **p**: P-value of effect estimate. **q**: Benjamini-Hochberg corrected P-value. **taxon:** microbial species tested.

**Table S11**. Results of association testing between milk CMV status and infant growth metrics at birth, 1 month, or 6 months of age. **est**: estimated effect of CMV+ vs. CMV- milk on infant trait. **se**: standard error of effect estimate. **p**: P-value of effect estimate. **trait:** infant growth metric and timepoint in the format trait_months; e.g. wlz_0 = weight-for-length at birth.

**Extended data file descriptions**

**milk_GeneExpr_TPM.txt**: gene expression levels estimated from RNA-sequencing of human milk, summarized as transcript per million (TPM). Rows are genes, columns are milk samples.

**milk_GeneExpr_counts.txt**: gene expression levels estimated from RNA-sequencing of human milk, summarized as counts. Rows are genes, columns are milk samples.

**metabolite_data.txt**: metabolite abundances from human milk. Rows are milk samples, columns are metabolites.

**infFecalMicrobiome_1month_taxonCLR.txt**: infant fecal microbiome taxon abundances (centered log ratio transformed) from 1 month of age. Rows are taxa, columns are infant fecal samples.

**infFecalMicrobiome_6month_taxonCLR.txt**: infant fecal microbiome taxon abundances (centered log ratio transformed) from 6 months of age. Rows are taxa, columns are infant fecal samples.
